# Effective host-directed therapy for tuberculosis by targeted depletion of myeloid-derived suppressor cells using a diphtheria toxin-based fusion protein

**DOI:** 10.1101/2020.12.10.420224

**Authors:** Sadiya Parveen, Shichun Lun, Michael E. Urbanowski, Mitchell Cardin, John R. Murphy, William R. Bishai

**Affiliations:** Department of Medicine, Johns Hopkins University School of Medicine, Baltimore, Maryland

**Author notes:** Co-corresponding authors William R. Bishai, Johns Hopkins School of Medicine, 1550 Orleans St, CRB2 Rm 108, Baltimore, MD 21287, John R. Murphy, Johns Hopkins School of Medicine, 1550 Orleans St, CRB2 Rm176, Baltimore MD 21287.

**Keywords:** Tuberculosis, Immunotherapy, Diphtheria fusion protein toxin, Host-directed therapy, MDSCs

## Abstract

Myeloid-derived suppressor cells (MDSCs) are present in elevated numbers in TB patients and have been found to be permissive for *Mycobacterium tuberculosis* (*Mtb*) proliferation. To determine whether depletion of MDSCs may improve host control of TB, we used a novel diphtheria toxin-based fusion protein known as DABIL-4 that targets and depletes IL-4-receptor positive cells. We show that DABIL-4 depletes both PMN-MDSCs and M-MDSC in the mouse TB model, and that it reduces the lung bacillary burden of *Mtb*. These results indicate that MDSC-depleting therapies targeting the IL4 receptor are beneficial in TB and offer an avenue towards host-directed TB therapy.

## INTRODUCTION

*Mycobacterium tuberculosis* (*Mtb*) is second only to COVID-19 in annual global mortality and caused the death of 1.7 million individuals in 2018 alone worldwide [1]. The treatment of TB is complicated by prolonged treatment time, patient compliance, drug-related toxicities, and the emergence of multi-(MDR) and extremely-drug resistance (XDR) strains. Thus, there is an urgent need to develop novel therapeutic approaches against *Mtb*. Host-directed therapies (HDTs) have the potential to reduce TB associated tissue damage and enhance host-mounted immunity, and they are insensitive to the antibiotic resistant state of the pathogen. In light of these attractive qualities, HDTs have been studied more intensively as an approach to adjunctive TB chemotherapy [2-5].

In recent years, MDSCs have emerged as a promising target for HDTs. MDSCs are a heterogenous population of progenitor-like immunocytes that suppress both innate and adaptive immunity in TB and many other disease settings. In murine TB models, MDSCs first appear at the site of infection and later disseminate to bronchi, pleural and peripheral blood, and similar patterns of expansion have also been noted in active TB patients [6, 7]. Of the two subsets of MDSCs, monocytic MDSCs (M-MDSCs) seem abundant in pleural effusions while polymorphonuclear-MDSCs (PMN-MDSCs) appear to be more prominent in bronchoalveolar lavage (BAL) of TB patients [7, 8]. Functionally, MDSCs suppress T-cell proliferation, dampen T-cell activation, and show no cytotoxic activity against mycobacteria [9-11]. Additionally, M-MDSCs also serve as safe harbor for *Mtb* replication owing to their unique metabolic status driven by IL-4/IL4R dependent mechanisms [12-14].

IL-4 receptor upregulation on myeloid-derived suppressor cells and M2-macrophages is known to contribute to their immunosuppressive activity of blocking effector T-cell mediated immunity in both infectious diseases as well as many cancers [15, 16]. IL-4R targeted therapies have been shown to deplete MDSCs and confer therapeutic benefits in cancer [17]. Based on these observations, we hypothesized that targeted depletion of IL-4R-bearing MDSCs may not only deplete an important *Mtb* intracellular niche but also alleviate local immunosuppression in tuberculous granulomas.

Diphtheria toxin-based fusion proteins have been used for targeted, selective cell depletion in animal models and in humans [18]. Owing to the enzymatic killing mechanism of diphtheria toxin, these recombinant proteins kill cells at the picomolar level [19]; they also have short serum half-lives of only 1-2 hours [20]. Therefore, when given systemically they lead to a transient depletion of cells which express the highest levels of the targeted receptor. We recently reported the development of a diphtheria toxin based fusion protein s-DAB_1-389_-mIL-4 (referred to as DABIL-4 hereafter) which showed an IC_50_ of 2 nM against 4T1, a murine breast adenocarcinoma cell line. We demonstrated that DABIL-4 targets and depletes IL-4R^+^ murine breast cancer cells, as well as tumor-associated MDSCs in a murine 4T1 adenocarcinoma model resulting in reductions of tumor growth and metastases [21]. Given its potency and immunotherapeutic role in cancer, we sought to determine whether DABIL-4-mediated MDSC depletion may play a beneficial role during tuberculosis. In the present study, we evaluated DABIL-4 in the murine model of TB and found that it selectively depletes IL-4R^+^ MDSCs and macrophages in both the lung and spleen and that this was accompanied by a concomitant reduction in the burden of *Mtb* in the lungs.

## RESULTS

### DABIL-4 selectively targets IL-4R expressing macrophages in vivo

As shown in **Fig 1A**, DABIL-4 is a fusion protein toxin which consists of the diphtheria toxin catalytic and translocation domains genetically fused to murine IL-4 (mIL-4). A related diphtheria toxin fusion protein in which IL-2 serves as the targeting domain (DABIL-2) was FDA approved as the drug denileukin diftitox, and it has been shown to have immunotherapeutic effects via depletion of IL-2 receptor (CD25^+^)-bearing Treg cells [19, 20, 22]. Prior to evaluating the potential therapeutic effect of DABIL-4 in a murine model of tuberculosis, we assessed the safety and tolerability of the drug in uninfected 129S2 mice. We administered three different doses of the fusion toxin (2.5, 5 and 10 µg / mouse / day) intraperitoneally to mice. After 21 days, mice treated with 5 and 10 µg of DABIL-4 daily showed significant weight loss in a dose-dependent fashion. In contrast mice treated with 2.5 µg daily and the vehicle-control group did not demonstrate weight loss (**Fig 1B**). Despite early weight loss, mice receiving 5 or 10 µg daily began to gain weight such that by day 21 they were almost at control levels. Importantly, despite the heavy dosing regimen none of the mice died (data not shown). On day 21, the mice were sacrificed and spleen cells were characterized by flow cytometry. As shown in **Fig. 1C**, we observed a dose dependent reduction of IL-4R^+^ macrophages in mice that were treated with daily DABIL-4; at the highest dose of 10 µg daily IL-4R^+^ macrophages were depleted by 3.5-fold. These results indicate that DABIL-4 administration is reasonably well tolerated and effectively depletes IL-4R^+^ macrophages in vivo.

**Fig 1.**
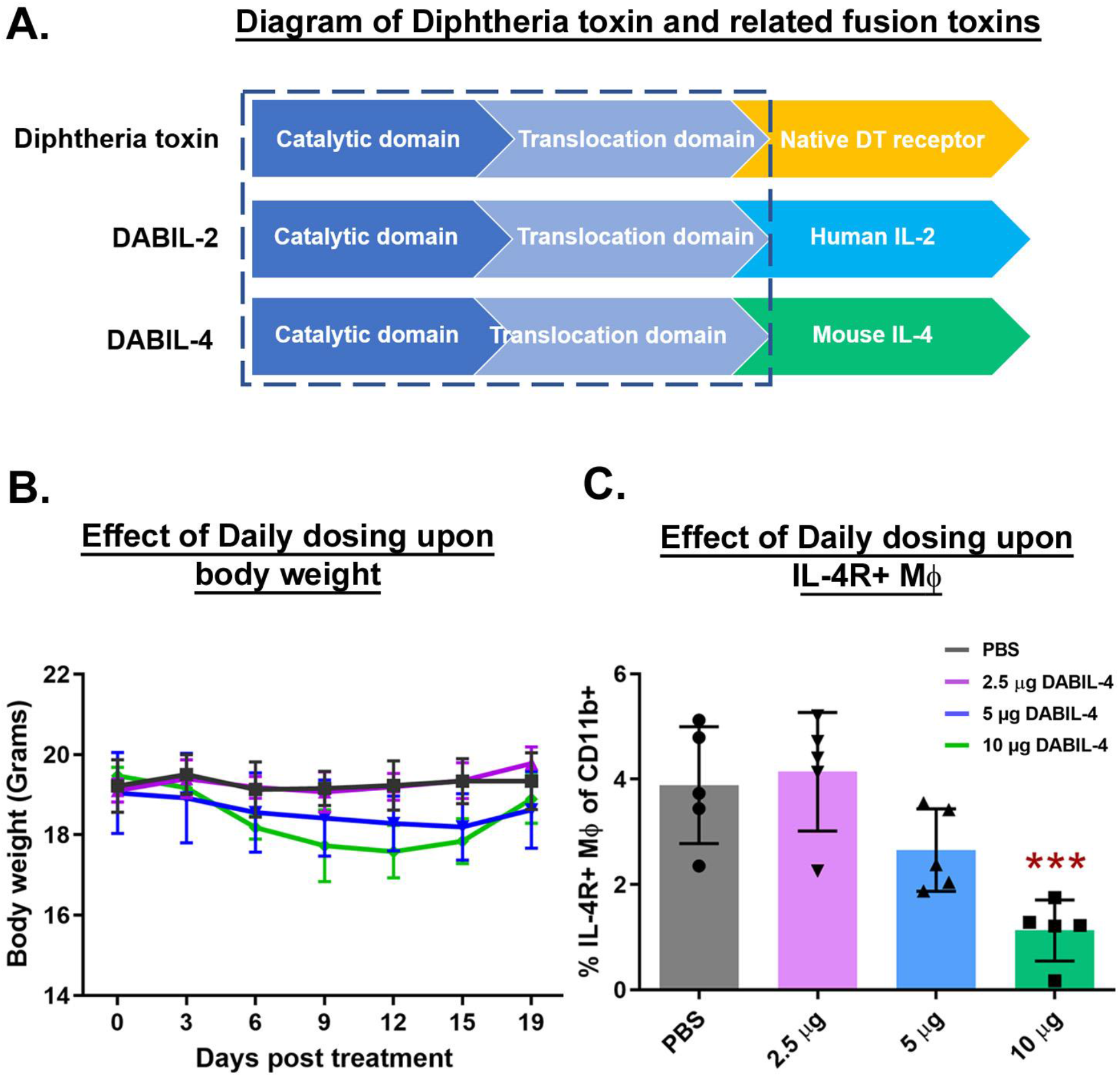
DABIL-4 safely targets IL-4R^+^ macrophages in vivo. **(A)** Schematic diagram showing domain structure of native diphtheria toxin and parent fusion toxin, DABIL-2 [19] compared to DABIL-4. The dotted blue box represents the domains critical for the cytotoxic activity of diphtheria toxin **(B)** Body weights of mice treated with indicated concentrations of DABIL-4 every day for 19 days. **(C)** Mice were treated with indicated concentrations of DABIL-4 every day and were sacrificed on day 21. Single cell suspensions of spleens from both PBS- and DABIL-4 treated groups were stained with specific antibodies and analyzed by multicolor flow cytometry (n=5). We found reduction in the population of IL-4R^+^ macrophages. Data are represented as mean ± SD and as percentage of CD11b^+^ population. Statistical significance between the groups was assessed by two-tailed unpaired student *t*-test considering an unequal distribution. \**p* < 0.05, \*\**p* < 0.01, \*\*\**p* < 0.001.

### DABIL-4 monotherapy inhibits mycobacterial replication in murine TB model

In contrast to wild-type mice, the C3HeB/FeJ (“Kramnik”) mouse strain is known to develop necrotic granulomatous lesions following *Mtb* infection similar to the pathology observed in human TB patients [23], and they also display elevated levels of MDSCs [13]. To evaluate the potential therapeutic efficacy of DABIL-4, we infected C3HeB/FeJ mice by the aerosol route and treated them with 10 µg DABIL-4 every third day starting on day 3 post-infection (**Fig. 2A**). The day 1 *Mtb* implantation in the lungs was 1.93 ± 0.14 log units (**Fig 2B**). After 21 days, mice from both the treated and untreated groups were sacrificed and lungs were collected. The DABIL-4 treated group had a quantitative lung bacterial burden that was 0.35 log units lower than that of the vehicle-treated control (*p* < 0.018) (**Fig 2C**). These data show that DABIL-4 monotherapy reduces *Mtb* proliferation in the lungs of infected mice.

**Fig 2.**
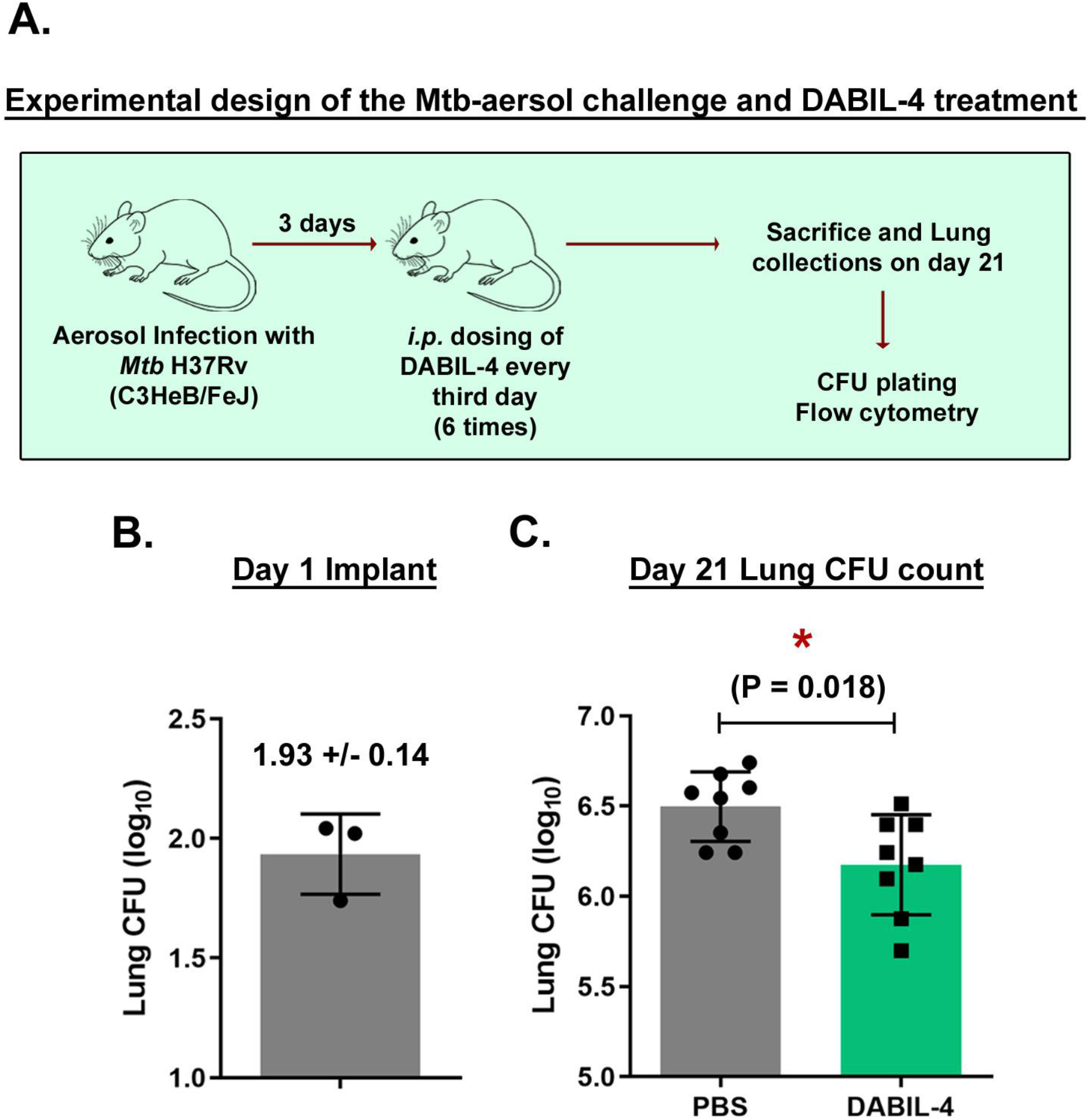
DABIL-4 treatment reduces lung bacillary burden in murine TB model. **(A)** Schematic of the experiment. C3HeB/FeJ mice (n=8) were aerosol infected with *Mtb* H37Rv. Mice were treated with 10 µg DABIL-4 on days 3, 6, 9, 12, 15, 18 post on days 3, 6, 9, 12, 15, 18 post aerosol infection. **(B)** On day 1 (n=3) and **(C)** Day 21 (n=8), lungs were collected from both PBS and DABIL-4 treated mice, homogenized, diluted and plated for quantification of bacillary burden in terms of CFU counts. Colonies were counted after 3-4 weeks. The CFU counts were transformed into log_10_ values and plotted using GraphPad Prism software. Statistical significance between the groups was assessed by two-tailed unpaired student *t*-test considering an unequal distribution. Data are shown as mean ± SD. \**p* < 0.05.

### DABIL-4 monotherapy reduces MDSCs in murine lungs during TB infection

Because MDSCs express high levels of the IL-4R (CD124) on their surface [24, 25], we sought to determine whether the therapeutic benefit of DABIL-4 was associated with MDSC depletion. To investigate the effect of DABIL-4 upon the host myeloid cell populations, we harvested lungs and spleens from both treated and untreated *Mtb*-infected C3HeB/FeJ mice on day 21 post treatment initiation (protocol shown in **Fig. 2A**), and single cell suspensions were characterized by multicolor flow cytometry. We first scored the effect upon myeloid cells and noted a significant increase in the total myeloid cell population (CD11b^+^ of all CD45^+^ cells) in both lungs and spleens (**Fig S1A** and **S1B**). Interestingly, despite an overall increase in myeloid cell abundance in the lungs, the population of IL-4R^+^ myeloid cells in lungs was reduced by 27% (*p* < 0.05) (**Fig 3A**), the observation is consistent with our breast cancer study [21]. We then characterized the impact of DABIL-4 administration specifically on MDSCs, a cell population known to drive immunosuppression in the granuloma. In DABIL-4 treated mice, there was a 36% reduction (p < 0.01) in the total MDSC population in lungs (**Fig 3B**). We also evaluated the impact on two distinct MDSCs subsets; polymorphonuclear MDSCs (PMN-MDSCs; CD11b^+^ Ly6G^+^ Ly6C^low^ CD124^+^) and monocytic-MDSCs (M-MDSCs; CD11b^+^ Ly6G^-^ Ly6C^high^ CD124^+^). We found close to a 30% reduction in the populations of both PMN-MDSCs and M-MDSCs (both *p* < 0.05) in the lung compartment (**Fig 3C** and **3D**). Interestingly, despite DABIL-4 depletion therapy, we saw a small increase in the M-MDSCs population in the spleen while the PMN-MDSC spleen population remained unchanged (data not shown) possibly reflecting accelerated replenishment of myeloid cells in the spleen. These results suggest that DABIL-4 effectively targets and depletes IL-4R^+^ immunosuppressive MDSCs in the lungs, most likely resulting in alleviation of immunosuppression and hence better host-mediated bacterial clearance in the lung.

**Fig 3.**
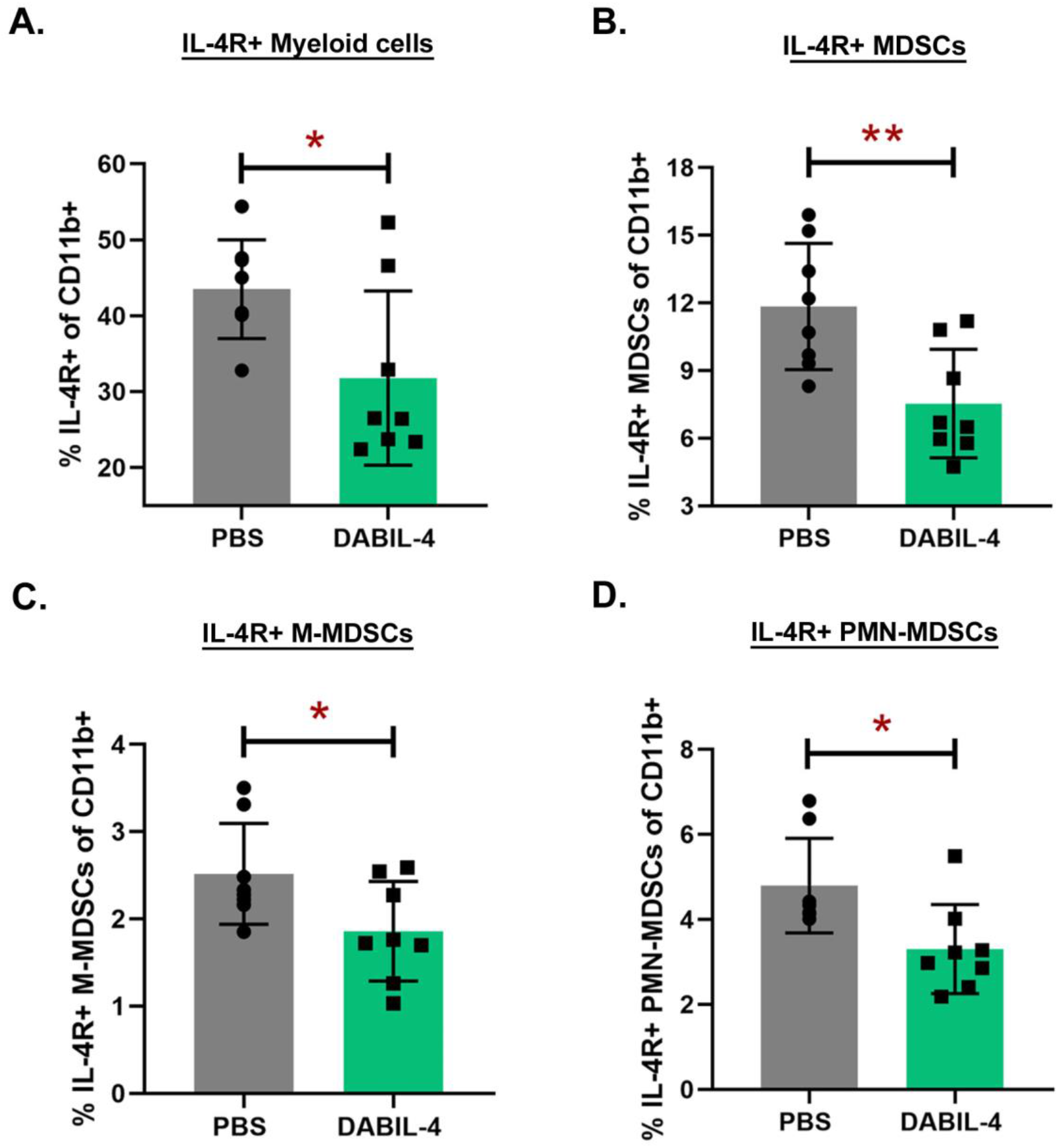
DABIL-4 administration depletes MDSCs population in lungs. As in **Fig 2A**, *Mtb*-infected C3HeB/FeJ mice were treated with DABIL-4 every third day beginning on day 3, and they were sacrificed on day 21. Single cell suspensions of lungs from both PBS- and DABIL-4 treated groups were stained with specific antibodies and analyzed by multicolor flow cytometry (n=6 to 8). We found differences in the population of **(A)** IL-4R^+^ of CD11b^+^ cells, **(B)** IL-4R^+^ MDSCs, **(C)** IL-4R^+^ M-MDSCs, **(D)** IL-4R^+^ PMN-MDSCs. Data are represented as mean ± SD and as percentage of CD11b^+^ population. Statistical significance between the groups was assessed by two-tailed unpaired student t-test considering an unequal distribution. \**p* < 0.05, \*\**p* < 0.01, ***P < 0.001.

### DABIL-4 monotherapy also targets other IL-4R-expressing macrophage populations in the murine TB model

We also investigated the impact of DABIL-4 on M2 macrophages, which not only serve as a safe harbor for *Mtb* replication and proliferation in the host, but are known also to suppress host immunity [26]. The presence of IL-4R on M2 macrophages specifically makes them susceptible to the DABIL-4 targeted toxin [27]. Using the 21-day infection and treatment protocol described in **Fig 2A**, we gated specifically on macrophages (F4/80^+^ CD124^+^ of all CD11b^+^ cells) and observed a 30% decline in the population of IL-4R^+^ macrophages (p < 0.001) (**Fig 4A**) harvested from DABIL-4 treated mice compared to the control group. Interestingly, DABIL-4 treatment did not perturb the population of IL-4R^+^ macrophages in spleen despite an overall increase in the macrophage population (**Fig S2**).

**Fig 4.**
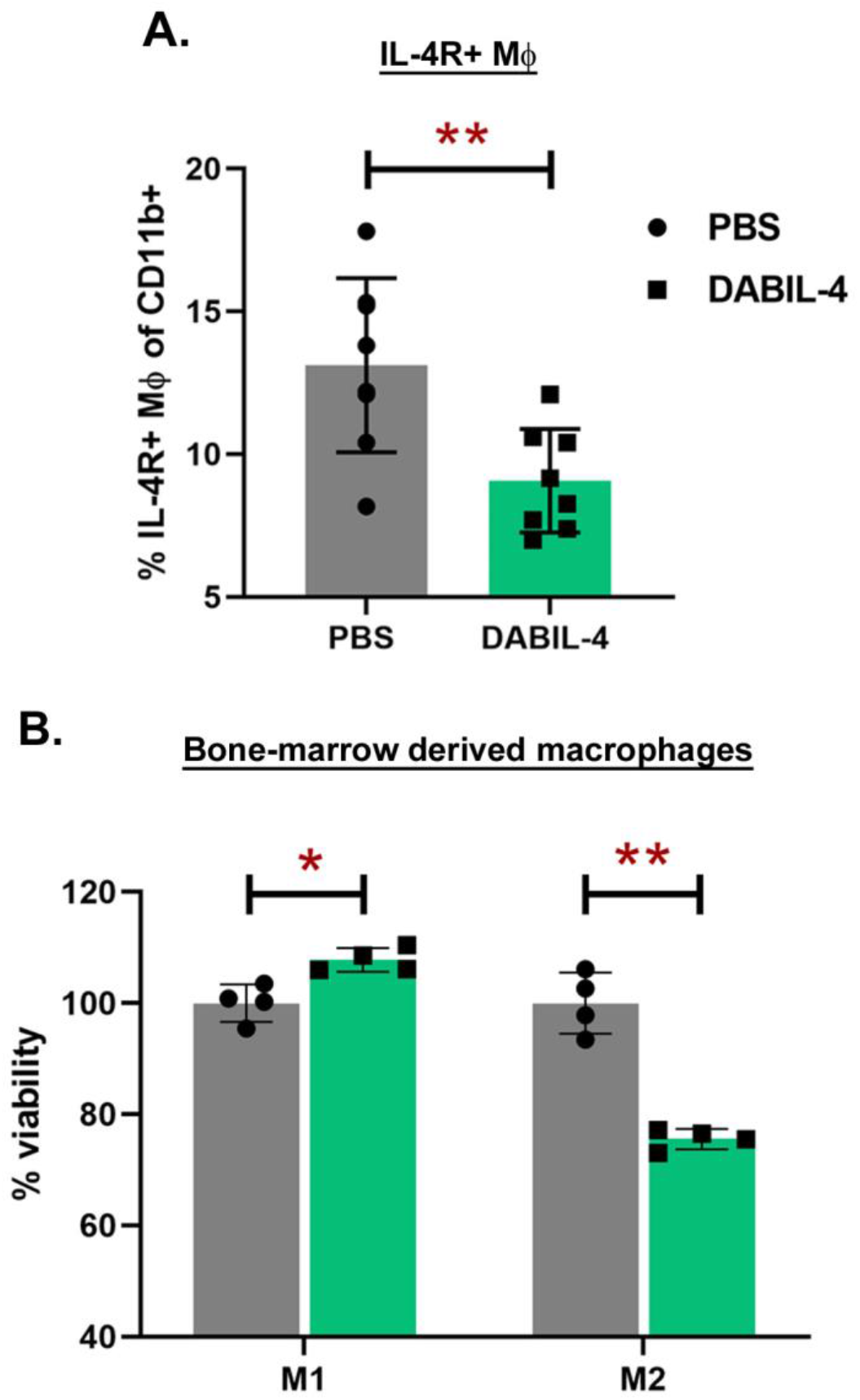
DABIL-4 administration depletes IL-4R^+^ macrophages in lungs. As shown in **Fig 2A**, *Mtb*-infected C3HeB/FeJ mice were treated with DABIL-4 every third day beginning on day 3, and they were sacrificed on day 21. Single cell suspensions of lungs from both PBS- and DABIL-4 treated groups were stained with specific antibodies and analyzed by multicolor flow cytometry (n=6 to 8). We found differences in the population of **(A)** IL-4R^+^ macrophages as percentage of CD11b^+^ population. **(B)** DABIL-4 treated-M2 polarized BMDMs showed a significant decline in viability. These BMDMs were polarized and treated with medium alone or medium containing 4 nM DABIL-4. The assay was performed as described in Materials and Methods. All values are expressed as Mean ± SD. Statistical significance between the groups was assessed by two-tailed unpaired student *t*-test considering an unequal distribution. \**p* < 0.05, \*\**p* < 0.01.

In order to investigate whether DABIL-4 selectively targets and depletes only M2-like macrophages, we isolated monocytes from the bone-marrow of healthy, uninfected mice and differentiated these cells into macrophages in presence of L929-conditioned media. We polarized these cells into M1- and M2-like macrophages, and then treated these populations with 4 nM DABIL-4 for 48 hours and scored their viability as described in the Materials and Methods section. We observed a significant decline in the viability of the DABIL-4 treated M2-macrophages (anti-inflammatory), whereas pro-inflammatory M1-like macrophages which do not express significant levels of IL-4R positivity showed no loss of viability and actually expanded moderately (**Fig 4B**). Based on these results we concluded that DABIL-4 selectively targets and depletes M2-like macrophages.

### DABIL-4 monotherapy modulates T cell populations during murine TB infection

In addition to myeloid cells, CD4^+^ T lymphocytes also are known to be key players in the establishment of protective immunity against tuberculosis in mice [28]. While active TB patients demonstrate an increased population of T-cells producing IL-4 in the periphery [29]. While the exact role played by these cells during TB infection is unclear, based on studies of humans with filarial-*Mtb* co-infections in which IL-4^+^ CD4^+^ T-cell populations are expanded, it has been proposed that they may potentially interfere with the Th1 cells expansion via Th1-Th2 crosstalk [30].

In light of these observations, we investigated the effect of DABIL-4 treatment upon both CD4^+^ IL-4R^+^ and CD8^+^ IL-4R^+^ T-cells in our murine TB model. Using the protocol outlined in **Fig. 2A**, we prepared cell suspensions from the lungs and spleens of *Mtb*-infected mice after 21 days of PBS or DABIL-4 treatment and analyzed them using multicolor flow cytometry. In lungs harvested from the DABIL-4 treated group, we observed a significant reduction in the population of IL-4R^+^ CD4^+^ IL-4R^+^ (**Fig 5A**) and CD8^+^ IL-4R^+^ T-cells (**Fig 5B**) both with *p* < 0.05. In spleen isolated from DABIL-4 treated mice, we did observe depletion of IL-4R^+^ CD4^+^ T-cells (**Fig 5C**) but IL-4R+ CD8+ T-cell population remain unchanged (data not shown). This loss of IL-4R expressing cells could be due to direct DABIL-4 targeting or indirect modulatory effects of MDSCs depletion.

**Fig 5.**
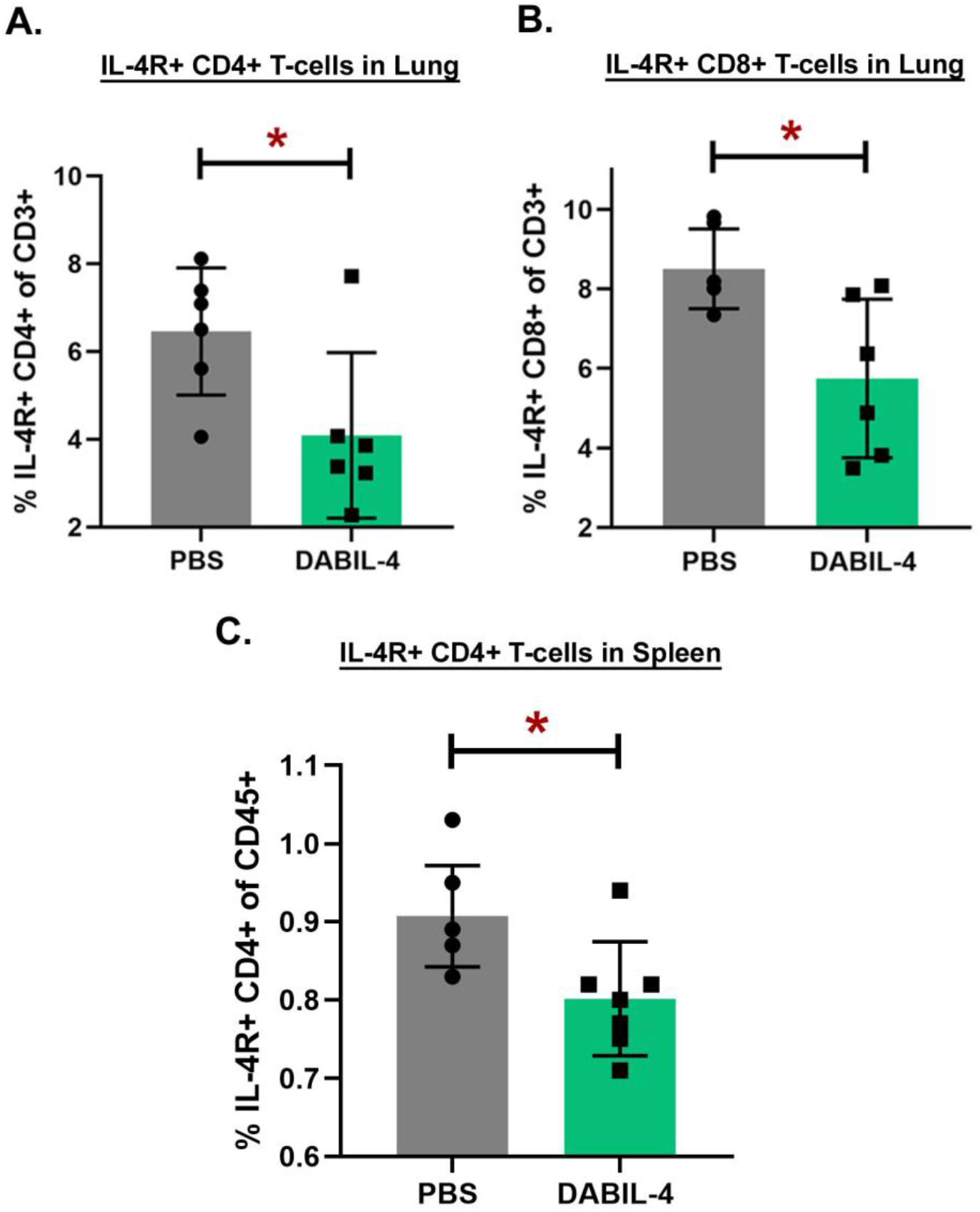
DABIL-4 administration depletes IL-4R^+^ lymphocytes while increasing the population of effector T-cells. As shown in **Fig 2A**, *Mtb*-infected C3HeB/FeJ mice were treated with DABIL-4 every third day beginning on day 3, and they were sacrificed on day 21. Single cell suspensions of lungs and spleens from both PBS- and DABIL-4 treated groups were stained with specific antibodies and analyzed by multicolor flow cytometry (n=6 to 8). We found differences in the population of **(A)** IL-4R^+^ CD4^+^ T-cells and **(B)** IL-4R^+^ CD8^+^ T-cells in the lung. We also saw decrease in **(C)** IL-4R+ CD4+ T-cells in spleen. Data are represented as mean ± SD and as percentage of CD3^+^ population except panel C which is represented as “percentage population of CD45+ cells. All values are expressed as Mean ± SD. Statistical significance between the groups was assessed by two-tailed unpaired student *t*-test considering an unequal distribution. \**p* < 0.05.

## DISCUSSION

Here, we report that targeting MDSCs and the IL-4/IL-4R axis in tuberculosis with DABIL-4 is an effective host directed immunotherapy in the mouse TB model. DABIL-4 is a diphtheria toxin-IL4 fusion protein related to the FDA-approved drug denileukin diftitox [31]. DABIL-4 kills IL-4R-bearing cells with an IC_50_ of 2 nM [21]. Related diphtheria toxin fusion proteins with similarly low IC_50_ values have been evaluated in humans and animals and are known to have short half-lives of ∼90 min [32]. Thus, parenteral administration of DABIL-4 is expected to deplete cells with high level expression of IL4 receptors such as MDSCs [24, 25] but in a transient manner owing to the short half-life. Our results in the mouse TB model show that DABIL-4 selectively depleted both IL-4R+ M- and PMN-MDSCs in mouse lungs and that this was accompanied by improved containment of *Mtb* proliferation in the lungs by CFU count determination. DAB-IL4 treatment also led to shifts in T cell populations including depletion of both IL-4R+ CD4+ and IL-4R+ CD8+ cells in *Mtb*-infected mice. The targeting of IL-4R+ T-cell could result from direct targeting by the toxin or an indirect consequence of MDSCs depletion and the resulting immunosuppression alleviation.

Lakkis et al. were the first to describe the genetic construction, purification and properties of DABIL-4. In their work, in vivo administration of DABIL-4 was found to be well tolerated and when administered subcutaneously in DBA/2 mice, treatment resulted in suppression of delayed-type hypersensitivity [33]. In light of that work suggesting that DABIL-4 was able to effectively modulate the immune responses, we constructed a gene fusion between the catalytic and translocation domains of diphtheria toxin and murine IL-4. In order to achieve expression and secretion of the fusion protein into the culture medium the structural gene also included the native diphtheria *tox* promoter, a mutant form of the *tox* operator, and the native *tox* signal sequence [19, 21]. Following expression and purification of DABIL-4, we confirmed and extended the earlier observations of Lakkis et al. by showing that the in vivo administration of the fusion protein toxin was well tolerated and resulted in the depletion of IL-4R+ positive myeloid cells in the spleen [21].

Following challenge of C3HeB/FeJ mice with *Mtb* H37Rv, DABIL-4 monotherapy reduced the burden of *Mtb* in the lungs by 0.35 log_10_ units (p = <0.018). The reduction of the *Mtb* bacillary burden may potentially be attributable to depletion of *Mtb*-harboring M2 macrophages as well as alleviation of immunosuppression due to MDSCs depletion in the infected lungs. M2 macrophages not only act as an intracellular niche for *Mtb* but also modulate granuloma formation and immune responses contributing to TB dissemination [34]. While there is an abundance of prior literature on the role of M2 macrophages, the potential immunosuppressive role of MDSCs has remained less well defined in TB pathogenesis (Parveen et al, Nature-Springer Press, unpublished). MDSCs are among the first cellular responders to the lungs following *Mtb* infection and have been associated with the granulomatous immune response in lung lesions and dampening of T-cell mediated anti-TB immunity [14, 35, 36]. However, the exact molecular mechanism by which MDSCs dampens T-cell immunity in TB remains to be deciphered.

The results reported here are consistent with earlier results using non-specific MDSC depletion strategies in the mouse TB model. In an earlier report, our group investigated the activity of tasquinimod, a quinoline-3-carboxamide analog that suppresses the local recruitment and function of MDSCs thorough its interaction with S100A9 [37], in *Mtb*-infected mice. We found that treatment with tasquinimod for 21 days significantly depleted the MDSC population and inhibited *Mtb* proliferation in both lungs and spleens of infected mice [38]. In another study by Knaul et al. used all-trans retinoic acid (ATRA), a retinoid known to reduce MDSC cells by promoting their differentiation to more mature myeloid cells via ERK1/2-dependent signaling and reduced levels of ROS [39, 40], was evaluated in the mouse TB model. Similar to tasquinimod, ATRA not only depleted MDSCs, but also reduced bacterial proliferation and improved lung pathology in mice [13]. In the present study, we have employed an IL-4 receptor targeted strategy using DABIL-4. Our results show that directed depletion of IL-4R expressing cells (MDSCs and M2 macrophages) produces comparable results to non-specific MDSC reducing agents such as tasquinimod and ATRA.

Interestingly, a previous study by Buccheri et al demonstrated that inhibition of IL-4/IL-4R axis with an IL-4 neutralizing antibody in BALB/c mice slowed TB progression [41]. While IL-4 depletion alone showed modest efficacy in reducing the bacterial count, combining it with an anti-mycobacterial α-crystallin mAb and IFNγ reduced the bacterial counts by 40-fold in the lungs at 3 weeks post infection. Moreover, the study found that BALB/c knockout mice lacking the IL-4 gene showed reduced *Mtb* proliferation compared to wild type [41]. Additionally, recombinant BALB/c with IL-4Rα^−/–^ knockout mice with concomitant IL-13 overexpression develop a progressive form of TB and demonstrate necrotic granulomas similar to those seen in humans [42]. These findings strongly support an important role of IL-4/IL-13 axis in TB pathogenesis, and suggest it may be an attractive target for the development of novel host-targeted therapies.

In conclusion, this proof of concept study conducted in the murine model of pulmonary TB demonstrates that the specific, directed depletion of IL-4R^+^ MDSCs using DABIL-4 is an effective host-directed therapy for TB in mice. Further studies are warranted to determine if DABIL-4 in combination with standard chemotherapy may help shorten the TB treatment duration. Additionally, the ability to specifically and transiently deplete IL-4R^+^ MDSCs with DABIL-4 offers a unique tool to study *Mtb* pathogenesis and to extend our understanding of the role of IL-4R/IL-4 axis in TB.

## MATERIALS AND METHODS

### Fermenter Protein purification of DABIL-4

DABIL-4 protein was purified using a protocol pioneered in our lab (Cheung et al 2019). Briefly, a *C. diphtheriae* non-lysogenic non-toxigenic C7s(-)*tox*-strain carrying pKN2.6Z-LC128 was grown in CY medium in a fermenter (Bioflo/Celligen 110). Supernatant was harvested as the culture reached OD ∼12-15, and concentrated by tangential flow filtration and a 30kD hollow fiber membrane (Spectrum). The flow concentrate was then diafiltered and adsorbed on to a HisTrap HP column (GE Healthcare). Bound protein was eluted with 50 mM NaH_2_PO_4_, 500 mM NaCl, 500 mM imidazole (pH 7.4). the protein was further concentrated using a 10 kDa Amicon Ultra-15 centrifugal unit (Millipore Sigma) and separated over a HiPrep 26/60 Sephacryl S100-HR sizing column (GE Healthcare Life Sciences). 5 ml fractions were collected and analyzed by SDS-PAGE. The protein concentration was then estimated and aliquots were stored at -80°C.

### Aerosol infection and CFU plating

We procured 129S2 mice from Charles River Laboratories (Wilmington, Massachusetts) and aerosol infected using *M. tuberculosis* H37Rv as previously described. All the protocols were approved by Johns Hopkins Institutional Animal Care and Use Committee. Post aerosol infection, three mice were sacrificed, their lungs were homogenized and plated for day 0 implant assessment. Rest of the mice were assigned into two groups. One group was treated with PBS while the other group received 10 µg/ml DABIL-4 every third day starting day 3 post infection as per the experimental design shown in Fig 2A. On day 21, lungs were harvested, homogenized, diluted and plated on selective middlebrook plates for the quantification of colony forming units (CFU).

### Bone-marrow derived macrophages (BMDMs) isolation and polarization

Murine bone marrow was isolated from 8-12-week-old female C57BL/6 mice. To differentiate into primary macrophages, bone marrow derived monocytes cells were incubated for 7 days in presence of RPMI medium-Glutamax (61870-036; Gibco) supplemented with 10% heat-inactivated FBS (16140071; Gibco) and antibiotics (penicillin-streptomycin solution; A5955; Sigma Aldrich) and 30% (vol/vol) L929 conditioned media. Mouse fibroblast L929 cells (ATCC CCL-1) were maintained in DMEM medium-Glutamax (10566-016; Gibco) supplemented with 10% FBS and antibiotics.

For polarization in 96 well plates, 500,000 cells per well were plated. After 3 hours and confirming the adherence, the cells were incubated with either M1-activation medium [RPMI160 medium with 20 ng/ml IFNγ (485-MI-100; R&D systems) and 10 ng/ml LPS (L2630; Sigma Aldrich)] or M2-activation medium [RPMI160 medium with 20 ng/ml IL-4 (404-ML-010; R&D systems) and 100 ng/ml MCSF (416-ML-010; R&D systems)] for 24 hours.

### Cytotoxicity assay with BMDMs and DABIL-4

For MTS assay, 50,000 BMDMs were plated per well in quadruplicate in 200 ul volume in 96-well plates, allowed to adhere, polarized and treated with medium containing 4 nM DABIL-4 fusion toxin or medium containing no drug. After 48 h incubation with the drug, [3-(4,5-dimethylthiazol-2-yl)-5-(3-carboxymethoxyphenyl)-2-(4-sulfophenyl)-2H-tetrazolium, inner salt; MTS reagent (Promega) was added to the individual wells. After 60 minutes, absorbance at 490 nM was recorded with the iMark Microplate Reader (Bio-Rad) to score cell viability. Wells with only medium was used to score the background absorbance.

### Multicolor Flow cytometry

For single cell suspension preparation, lungs were chopped into small pieces and digested with collagenase D and DNaseI at 37 C for 1 h while spleens were passed through 100 µm mesh filters. To remove red blood cells, cell suspension was treated with RBC lysis buffer as per the supplier’s instructions (Biolegend). Single-cell suspensions from lung and spleen were scored for viability using trypan blue staining. To prevent non-specific binding, one million cells were incubated first with purified anti-mouse CD16/32 antibody (Biolegend, Cat# 101320) in FACS buffer (eBiosciences, Cat# 422226) and then stained with flow antibodies (Biolegend unless otherwise indicated); (1) Lymphoid panel: APC/Cy7 CD45, BV785 CD3, BUV563 CD4 (BD), AF700 CD8a, BV711 CD25, BV650 CD11b, BV605 Ly6G (BD), PE/CY7 Ly6C, PE CD124, APC F4/80. Zombie-UV fixable viability dye was included to select for viable cells. Post surface staining, cells were fixed in fixation buffer and washed twice with FACS buffer. The samples were recorded on LSRFortessa™ X-20 Cell Analyzer (BD) and using FlowJo v.10 software (Tree Star) was used for data analysis. FSC-W and SSC-H gates were designed to exclude debris and aggregates while FSC-A versus SSC-A gates were used to exclude duplets. Inside this latter gate, cells with positive Zombie-UV staining were excluded to remove dead cells from the analysis. All reagents were from Biolegend unless otherwise indicated.

### Statistical Analysis

All CFU counts were log10 transformed before analysis. Mean CFU counts were compared using two-tailed Student *t* tests. All measures of statistical variation are expressed as ± standard deviation of the mean (SEM).

## AUTHOR CONTRIBUTIONS

SP, JRM, and WRB conceptualized the study and designed the research approach; SP, SL, MEU and MC performed research; JRM contributed new reagents/analytic tools; SP, JRM, WRB analyzed data; and SP, JRM, and WRB wrote the paper, SP, SL, MEU, MC, JRM, DS, WRB critically reviewed the manuscript.

## ACKNOWLEDGEMENTS

We gratefully acknowledge the support of NIH grants AI 130595 and AI 1522688, Tedco awards 2016-MII-3464 and 2019-MII-518, and awards from the Abell Foundation and the Maryland Cigarette Restitution fund. We thank Dr. Geetha Srikrishna for editorial assistance and Stefanie Krug for helpful suggestions.

---

**Fig S1.**
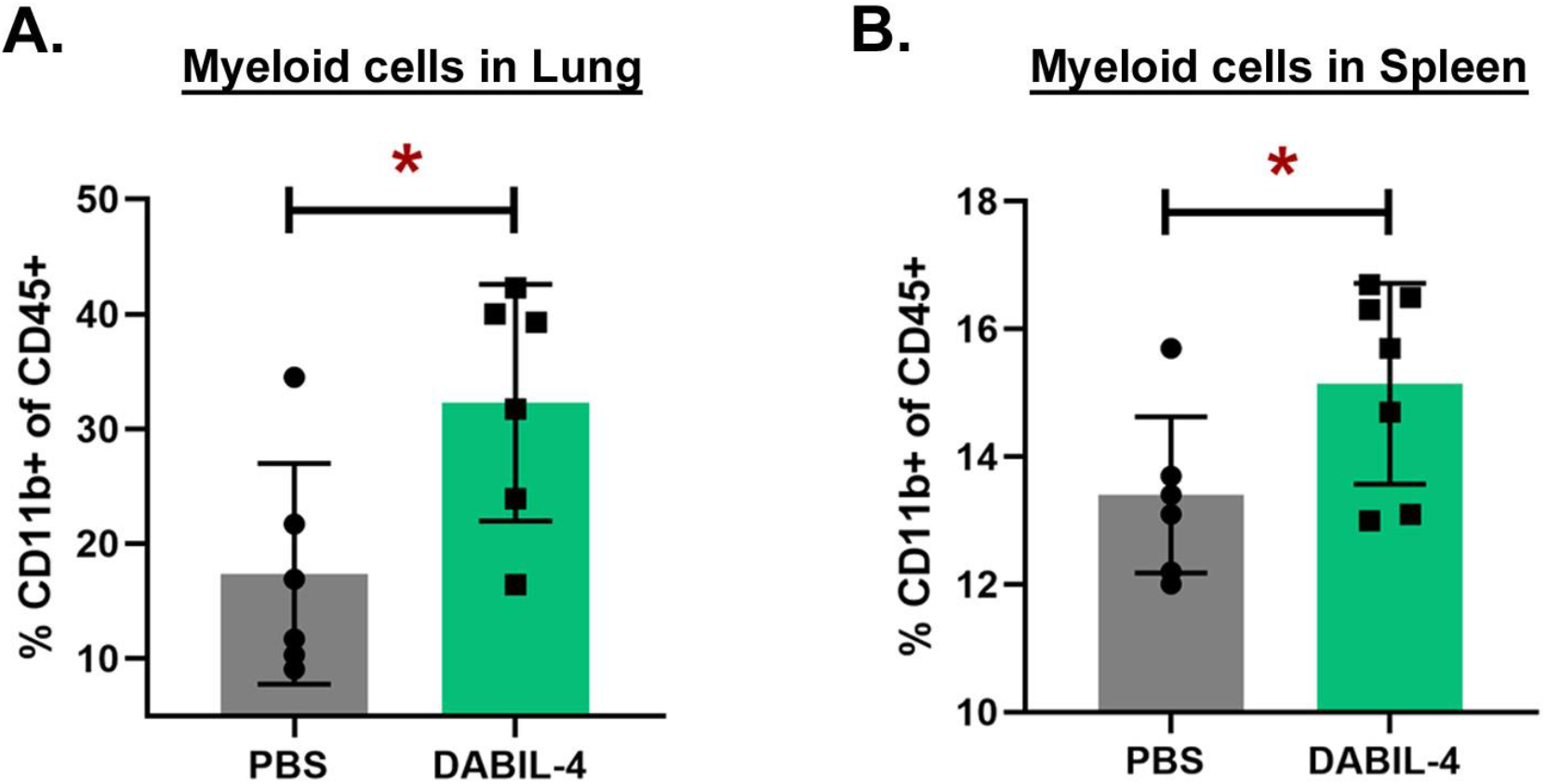
DABIL-4 treatment increases myeloid cell populations in the lung and spleen. On day 21, single cell suspensions of lungs and spleens from both PBS- and DABIL-4 treated groups were stained with specific antibodies and analyzed by multicolor flow cytometry (n=6 to 8). We found an increase in the population of CD11b^+^ expressed as percentage of CD45^+^ cells. Statistical significance between the groups was assessed by two-tailed unpaired student *t*-test considering an unequal distribution. Data are represented as mean ± SD. \**p* < 0.05.

**Fig S2.**
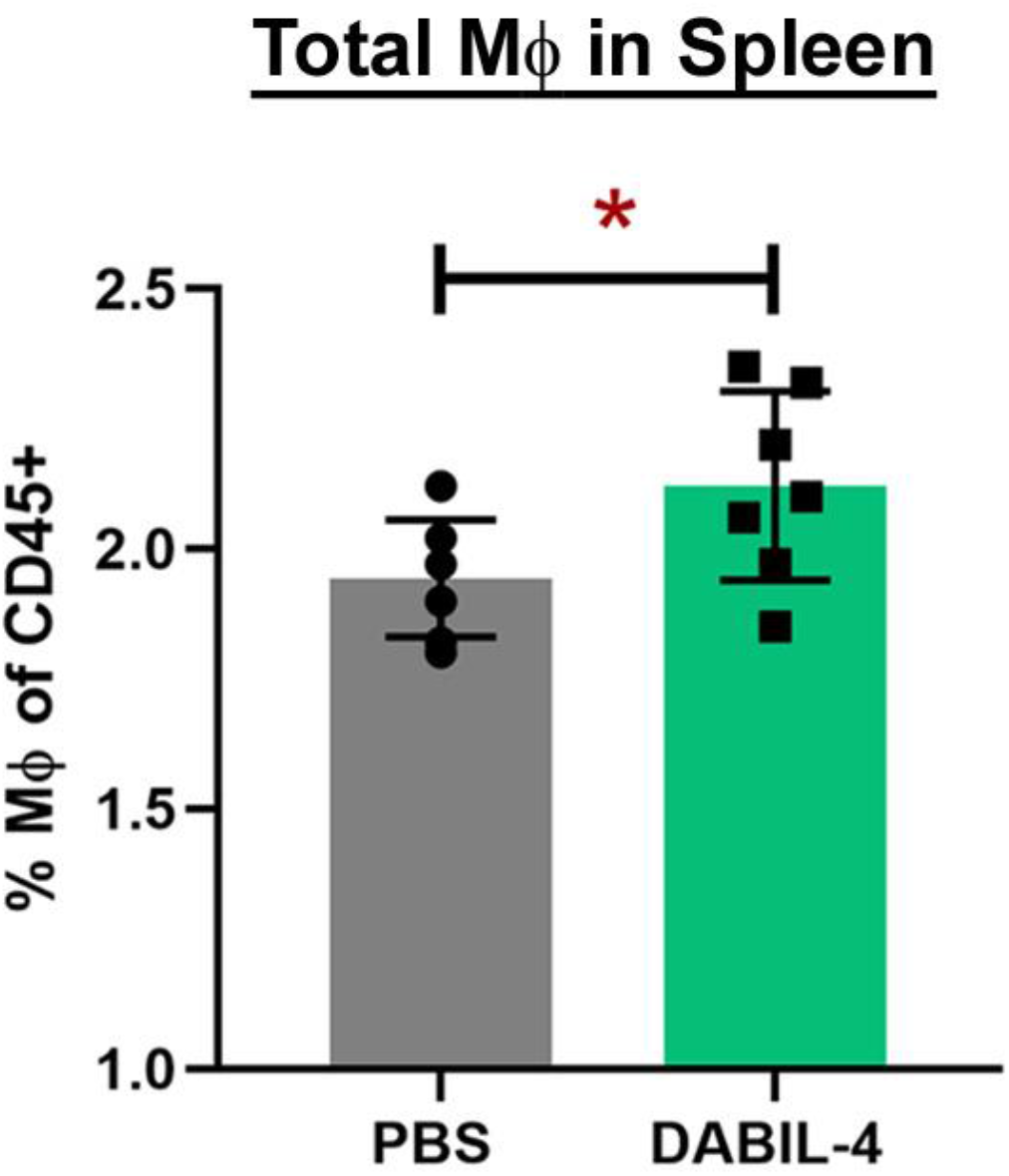
DABIL-4 treatment increases macrophage populations in spleen. On day 21, single cell suspensions of spleens from both PBS- and DABIL-4 treated groups were stained with specific antibodies and analyzed by multicolor flow cytometry (n=7). We found an increase in the population of macrophages (CD11b^+^ F4/80^+^) expressed as percentage of CD45^+^ cells. Statistical significance between the groups was assessed by two-tailed unpaired student *t*-test considering an unequal distribution. Data are represented as mean ± SD. \**p* < 0.05.

## BIBLIOGRAPHY

1. Geneva-World-Health-Organization. Global tuberculosis report 2020. Licence: CC BY-NC-SA 30 IGO 2020.

2. Tiberi S, du Plessis N, Walzl G, et al. Tuberculosis: progress and advances in development of new drugs, treatment regimens, and host-directed therapies. Lancet Infect Dis 2018; 18:e183–e98.

3. Kaufmann SHE, Dorhoi A, Hotchkiss RS, Bartenschlager R. Host-directed therapies for bacterial and viral infections. Nature reviews Drug discovery 2018; 17:35–56.

4. Young C, Walzl G, Du Plessis N. Therapeutic host-directed strategies to improve outcome in tuberculosis. Mucosal Immunol 2020; 13:190–204.

5. Abreu R, Giri P, Quinn F. Host-Pathogen Interaction as a Novel Target for Host-Directed Therapies in Tuberculosis. Frontiers in immunology 2020; 11.

6. Du Plessis N, Jacobs R, Gutschmidt A, et al. Phenotypically resembling myeloid derived suppressor cells are increased in children with HIV and exposed/infected with Mycobacterium tuberculosis. European journal of immunology 2017; 47:107–18.

7. Tsiganov EN, Verbina EM, Radaeva TV, et al. Gr-1dimCD11b+ immature myeloid-derived suppressor cells but not neutrophils are markers of lethal tuberculosis infection in mice. J Immunol 2014; 192:4718–27.

8. El Daker S, Sacchi A, Tempestilli M, et al. Granulocytic myeloid derived suppressor cells expansion during active pulmonary tuberculosis is associated with high nitric oxide plasma level. PloS one 2015; 10:e0123772.

9. Wang Z, Jiang J, Li Z, Zhang J, Wang H, Qin Z. A myeloid cell population induced by Freund adjuvant suppresses T-cell-mediated antitumor immunity. Journal of immunotherapy (Hagerstown, Md : 1997) 2010; 33:167–77.

10. Kato K, Yamamoto K. Suppression of BCG cell wall-induced delayed-type hypersensitivity by BCG pre-treatment. II. Induction of suppressor T cells by heat-killed BCG injection. Immunology 1982; 45:655–61.

11. Klimpel GR, Okada M, Henney CS. Inhibition of in vitro cytotoxic responses by BCG-induced macrophage-like suppressor cells. II. Suppression occurs at the level of a “helper” T cell. Journal of immunology (Baltimore, Md : 1950) 1979; 123:350–7.

12. Hossain F, Al-Khami AA, Wyczechowska D, et al. Inhibition of Fatty Acid Oxidation Modulates Immunosuppressive Functions of Myeloid-Derived Suppressor Cells and Enhances Cancer Therapies. Cancer Immunol Res 2015; 3:1236–47.

13. Knaul JK, Jorg S, Oberbeck-Mueller D, et al. Lung-residing myeloid-derived suppressors display dual functionality in murine pulmonary tuberculosis. Am J Respir Crit Care Med 2014; 190:1053–66.

14. Agrawal N, Streata I, Pei G, et al. Human Monocytic Suppressive Cells Promote Replication of Mycobacterium tuberculosis and Alter Stability of in vitro Generated Granulomas. Frontiers in immunology 2018; 9:2417.

15. Suzuki A, Leland P, Joshi BH, Puri RK. Targeting of IL-4 and IL-13 receptors for cancer therapy. Cytokine 2015; 75:79–88.

16. Ost M, Singh A, Peschel A, Mehling R, Rieber N, Hartl D. Myeloid-Derived Suppressor Cells in Bacterial Infections. Front Cell Infect Microbiol 2016; 6:37.

17. Roth F, De La Fuente AC, Vella JL, Zoso A, Inverardi L, Serafini P. Aptamer-mediated blockade of IL4Ralpha triggers apoptosis of MDSCs and limits tumor progression. Cancer research 2012; 72:1373–83.

18. Parveen S, Bishai WR, Murphy JR. Corynebacterium diphtheriae: Diphtheria Toxin, the tox Operon, and Its Regulation by Fe2(+) Activation of apo-DtxR. Microbiol Spectr 2019; 7.

19. Cheung LS, Fu J, Kumar P, et al. Second-generation IL-2 receptor-targeted diphtheria fusion toxin exhibits antitumor activity and synergy with anti-PD-1 in melanoma. Proceedings of the National Academy of Sciences of the United States of America 2019; 116:3100–5.

20. Manoukian G, Hagemeister F. Denileukin diftitox: a novel immunotoxin. Expert Opin Biol Ther 2009; 9:1445–51.

21. Parveen S, Siddharth S, Cheung LS, et al. IL-4 receptor targeting as an effective immunotherapy against triple-negative breast cancer. bioRxiv 2020:2020.08.05.238824.

22. Gupta S, Cheung L, Pokkali S, et al. Suppressor Cell-Depleting Immunotherapy With Denileukin Diftitox is an Effective Host-Directed Therapy for Tuberculosis. J Infect Dis 2017; 215:1883–7.

23. Pan H, Yan BS, Rojas M, et al. Ipr1 gene mediates innate immunity to tuberculosis. Nature 2005; 434:767–72.

24. Mandruzzato S, Solito S, Falisi E, et al. IL4Ralpha+ myeloid-derived suppressor cell expansion in cancer patients. Journal of immunology (Baltimore, Md : 1950) 2009; 182:6562–8.

25. Gallina G, Dolcetti L, Serafini P, et al. Tumors induce a subset of inflammatory monocytes with immunosuppressive activity on CD8+ T cells. The Journal of clinical investigation 2006; 116:2777–90.

26. Cohen SB, Gern BH, Delahaye JL, et al. Alveolar Macrophages Provide an Early Mycobacterium tuberculosis Niche and Initiate Dissemination. Cell Host Microbe 2018; 24:439-46.e4.

27. Feldman GM, Ruhl S, Bickel M, Finbloom DS, Pluznik DH. Regulation of interleukin-4 receptors on murine myeloid progenitor cells by interleukin-6. Blood 1991; 78:1678–84.

28. Yao S, Huang D, Chen CY, Halliday L, Wang RC, Chen ZW. CD4+ T cells contain early extrapulmonary tuberculosis (TB) dissemination and rapid TB progression and sustain multieffector functions of CD8+ T and CD3-lymphocytes: mechanisms of CD4+ T cell immunity. Journal of immunology (Baltimore, Md : 1950) 2014; 192:2120–32.

29. van Crevel R, Karyadi E, Preyers F, et al. Increased production of interleukin 4 by CD4+ and CD8+ T cells from patients with tuberculosis is related to the presence of pulmonary cavities. J Infect Dis 2000; 181:1194–7.

30. Chatterjee S, Clark CE, Lugli E, Roederer M, Nutman TB. Filarial infection modulates the immune response to Mycobacterium tuberculosis through expansion of CD4+ IL-4 memory T cells. Journal of immunology (Baltimore, Md : 1950) 2015; 194:2706–14.

31. Kumar P, Kumar A, Parveen S, Murphy JR, Bishai W. Recent advances with Treg depleting fusion protein toxins for cancer immunotherapy. Immunotherapy 2019; 11:1117–28.

32. Eklund JW, Kuzel TM. Denileukin diftitox: a concise clinical review. Expert Review of Anticancer Therapy 2005; 5:33–8.

33. Lakkis F, Steele A, Pacheco-Silva A, Rubin-Kelley V, Strom TB, Murphy JR. Interleukin 4 receptor targeted cytotoxicity: genetic construction and in vivo immunosuppressive activity of a diphtheria toxin-related murine interleukin 4 fusion protein. European journal of immunology 1991; 21:2253–8.

34. Shim D, Kim H, Shin SJ. Mycobacterium tuberculosis Infection-Driven Foamy Macrophages and Their Implications in Tuberculosis Control as Targets for Host-Directed Therapy. Frontiers in immunology 2020; 11.

35. Ostrand-Rosenberg S, Sinha P. Myeloid-derived suppressor cells: linking inflammation and cancer. Journal of immunology (Baltimore, Md : 1950) 2009; 182:4499–506.

36. du Plessis N, Kotze LA, Leukes V, Walzl G. Translational Potential of Therapeutics Targeting Regulatory Myeloid Cells in Tuberculosis. Front Cell Infect Microbiol 2018; 8:332.

37. Yoshioka Y, Mizutani T, Mizuta S, et al. Neutrophils and the S100A9 protein critically regulate granuloma formation. Blood Adv 2016; 1:184–92.

38. Gupta S, Krug S, Pokkali S, et al. Pharmacologic Exhaustion of Suppressor Cells with Tasquinimod Enhances Bacterial Clearance during Tuberculosis. Am J Respir Crit Care Med 2019; 199:386–9.

39. Nefedova Y, Fishman M, Sherman S, Wang X, Beg AA, Gabrilovich DI. Mechanism of all-trans retinoic acid effect on tumor-associated myeloid-derived suppressor cells. Cancer research 2007; 67:11021–8.

40. Draghiciu O, Lubbers J, Nijman HW, Daemen T. Myeloid derived suppressor cells-An overview of combat strategies to increase immunotherapy efficacy. Oncoimmunology 2015; 4:e954829.

41. Buccheri S, Reljic R, Caccamo N, et al. IL-4 depletion enhances host resistance and passive IgA protection against tuberculosis infection in BALB/c mice. European journal of immunology 2007; 37:729–37.

42. Heitmann L, Abad Dar M, Schreiber T, et al. The IL-13/IL-4Ralpha axis is involved in tuberculosis-associated pathology. J Pathol 2014; 234:338–50.

